# Evaluating Clustering and Imputation Approaches for Multivariate Longitudinal Clinical Data Using a Real-World eCDF-Copula Resampling Framework

**DOI:** 10.64898/2026.09.24.753982

**Authors:** Hanchang Cai, Ke Xu, Sarrah Osmundson, Olusola Ajilore, Carmen Andreescu, Warren D Taylor, Jinyuan Liu, Xinlian Zhang

**Author notes:** Corresponding author: Jinyuan Liu, Ph.D., 2525 West End Avenue, Suite 1100, Room 1137, Nashville, TN 37203, USA. Corresponding author: Xinlian Zhang, Ph.D., 9500 Gilman Drive, La Jolla, CA 92093, USA.

## Abstract

The rapid growth of technologies such as wearable devices and high-throughput sequencing has generated unprecedented volumes of longitudinal data, often with multiple correlated features. This trend highlights the need for robust clustering methods tailored to multi-feature longitudinal datasets. Although previous reviews have benchmarked clustering methods in this context, most evaluations relied heavily on simulated and overly simplified datasets, which substantially limit their interpretability and applicability to real-world data. Missing data is also ubiquitous in longitudinal data, yet understudied in the context of clustering. To address these gaps, we systematically evaluate clustering methods across three distinct types of real-world longitudinal datasets: (1) a high-dimensional, low-timepoint metagenomic study, (2) a low-dimensional, high-timepoint multi-site repeated-measure study, and (3) a high-dimensional, high-timepoint repeated-measure study. We further investigate how missing data and different imputation strategies influence clustering performance. In addition, we introduce a novel resampling framework based on the empirical cumulative distribution function (eCDF) and a copula. This framework enables us to simulate multi-feature longitudinal data that preserves the correlation structure of real datasets, while also allowing for controlled variations such as sample size. Our simulation framework naturally accommodates missing data, thereby providing a flexible tool for benchmarking and comparing clustering methods in more realistic longitudinal settings.

## 1 Introduction

Longitudinal multivariate data are becoming increasingly common across diverse domains, ranging from digital health technologies, such as wearable devices [1] and ecological momentary assessments (EMA) [2], to biomedical fields including imaging, genomics [3], and metagenomics [4, 5]. These datasets capture dynamic processes with multiple correlated feature trajectories and offer rich opportunities to study population heterogeneity. However, their complexity arises from high dimensionality, temporal correlation, and irregular sampling, with ubiquitous missingness posing unique methodological challenges.

Clustering analysis is a fundamental tool for uncovering latent subgroups within heterogeneous populations [6]. By partitioning individuals based on their joint longitudinal trajectories, clustering facilitates the discovery of disease subtypes, behavioral phenotypes, and other meaningful patterns that may inform clinical interventions or scientific hypotheses [7, 8]. The clustering of single longitudinal features has been extensively studied, where existing approaches are broadly divided into model-based methods (e.g., latent growth and growth mixture models [9, 10]) and nonparametric algorithms (e.g., k-means [11] and traj [12] methods). Recent studies demonstrate that `kml` [13] method based on k-means algorithm generally outperforms alternative approaches in terms of accuracy, separation, and stability, particularly when sufficient time points are available [14]. However, extending these approaches to jointly clustering multi-feature trajectories requires careful consideration of both statistical modeling and computational scalability.

A recent tutorial by Lu et al. (2023) [15] provided a comprehensive overview of existing clustering approaches for multiple longitudinal features, categorizing them into model-based methods (e.g., `gbmt` [16], `flexmix` [17–20], `lcmm` [9], `mixAK` [21, 22], `BCClong` [23]) and algorithm-based methods (e.g., `kml3d` [24, 25]). Their study highlighted key methodological differences, such as the treatment of random effects and assumptions about feature dependencies. They also illustrated these methods using simulated and empirical examples. However, as the authors acknowledged, their benchmark was largely model-driven and limited in its coverage of real-world heterogeneity. Further, performance under varying data structures and missing data mechanisms was not systematically assessed.

Beyond this, recent applied studies have started to demonstrate challenges of clustering multivariate longitudinal data in real-world contexts. For example, Stoitsas et al. (2022) [26] applied `kml3d`, `HDclassif`, and `DeepGMM` to trauma patient recovery trajectories and showed that different algorithms can yield clusters with varying levels of clinical interpretability, especially when missing data are imputed using chained equations. Similarly, Wenzel et al. (2024) [27] clustered 7-day EMA time-series of psychotic symptoms using dynamic time warping (DTW), identifying clinically meaningful high-symptom versus low-symptom clusters not captured by diagnostic categories. In microbiome research, Kodikara et al. (2022) [5] systematically compared clustering algorithms on longitudinal metagenomic data, showing that k-medoids and DTW-based clustering can reveal biologically relevant taxa groups, but performance deteriorates with increasing noise. Collectively, these works highlight the importance of benchmarking clustering strategies under diverse real-world conditions, while also exposing limitations such as sensitivity to imputation choices and data preprocessing.

These gaps highlight several unresolved issues. First, most existing evaluations focus on either single-feature or low-dimensional settings, and it is not clear how clustering algorithms generalize to richer, multi-feature longitudinal data. Second, while missingness is ubiquitous in longitudinal research, the influence of different imputation strategies on clustering performance has received limited attention. Third, simulation frameworks used for benchmarking often rely on strong parametric assumptions that may not capture the oftentimes skewed, multimodal, or bounded distributions observed in real-world clinical and behavioral datasets. In response, our study aims to improve upon real-world evaluation, add a missing-data assessment, and contribute a new nonparametric simulation framework. In particular, we aim to:

1. Benchmark clustering methods on three distinct real-life longitudinal scenarios: high-dimensional/low-timepoint, low-dimensional/high-timepoint, and high-dimensional/high-timepoint data.
2. Assess how different imputation strategies influence clustering validity and the selection of the number of clusters.
3. Propose an eCDF-copula-based resampling method that preserves the correlation structures of empirical datasets while allowing flexible specification of sample size or time points. We will generate synthetic data using this additional nonparametric approach to resemble their real-world patterns, which will facilitate fair cross-scenario comparisons and enable robust benchmarking in the presence of missingness.

## 2 Clustering Algorithms

We first review the clustering algorithms considered in our study. Let *y*_*ijk*_ denote the observation of individual *i* (*i* = 1, …, *n*), on feature *j* (*j* = 1, …, *p*), at time point *k* (*k* = 1, …, *t*). After imputation, both *p* and *t* are constant across individuals. The cluster membership of individual *i* is denoted by *C*_*i*_ ∈ {1, …, *G*}, where *G* is the number of clusters, with cluster weights *π*_*g*_ such that 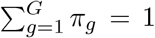. For notation, **y**_*ij*_ = (*y*_*ij*1_, …, *y*_*ijt*_)^*⊤*^ denotes the vector of feature *j* for subject *i* over time, and **Y**_*i*_ ∈ ℝ^*p×t*^ represents the subject-specific multi-feature matrix over time. Let **x** = (*x*_1_, …, *x*_*t*_)^*⊤*^ denote the corresponding measurement times shared across all features, where each *x*_*k*_ represents the time point (or visit) at which *y*_*ijk*_ is observed.

### 2.1 Group-based trajectory models

Group-based trajectory models (GBTM) [28, 29] assume that the study population is heterogeneous and can be described as a finite mixture of latent trajectory groups. The general finite mixture model can be written as

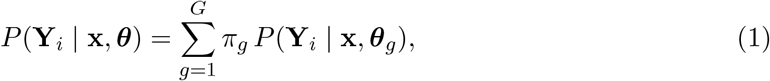

where *π*_*g*_ denotes the mixing proportion (or cluster weights) and ***θ***_*g*_ represents the class-specific parameters.

Conditional on class membership (*C*_*i*_ = *g*), features and time points are assumed independent. Each feature trajectory is modeled as a polynomial function of time with Gaussian errors, where the joint multi-feature probability becomes:

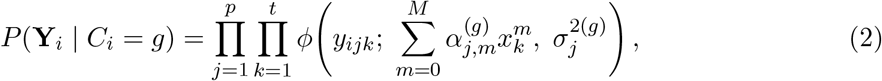

where *ϕ*(; *µ, σ*^2^) denotes the Gaussian density with mean *µ* and variance *σ*^2^, and *M* is the degree of the polynomial. Each individual belongs to one latent class *g*, within which trajectories are parameterized by 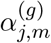.

Then the posterior membership probability of subject *i* can be readily calculated:

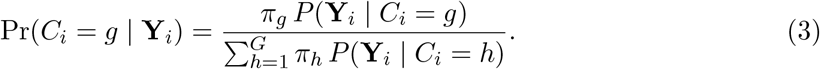

In practice, subjects are assigned to the class with the largest posterior probability. Both the R package `gbmt` and `flexmix` implement the GBTM framework.

### 2.2 Latent class mixed-effects model

Latent class mixed-effects models (LCMM) [9] extend GBTM by incorporating random effects to account for the correlation between repeated measurements. In the general setting, the conditional distribution of the outcome belongs to the exponential family F with a link function *h*(·). For subject *i*, feature *j*, and time point *k*, the model is specified as

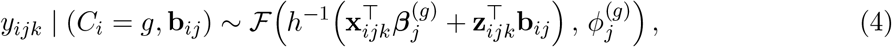

where **x**_*ijk*_ and **z**_*ijk*_ denote covariates for fixed and random effects, respectively. The random effect is assumed to follow a multivariate normal (MVN) distribution

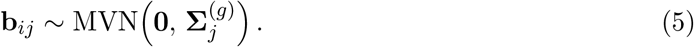

Each latent class *g* is characterized by its own fixed effects 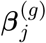 and random-effects covariance structure 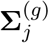. In principle, LCMM can accommodate different outcome types from the exponential family and is implemented using `lcmm` package in R. In our application, we focused on Gaussian outcomes for computational ease [30]. To avoid over-parameterization, we included only a random intercept, resulting in class-specific intercepts but shared random slopes across classes.

### 2.3 Bayesian mixture model

In contrast to LCMM, which allows both fixed and random effects to vary between classes, the Bayesian mixture model (BMM) [31] clusters subjects by modeling the random effects as coming from a finite mixture distribution while sharing common fixed effects across clusters. BMM further adopts a fully Bayesian framework with posterior inference via Markov chain Monte Carlo (MCMC) sampling. Let 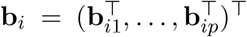 denote the concatenated random-effect vector of dimension *q* for subject *i*, where 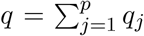 and *q*_*j*_ is the number of random-effect terms specified for feature *j*. For example, when only random intercepts are included, *q* = *p*; if both random intercepts and random slopes with respect to time are included, *q* = 2*p*.

For subject *i* and feature *j* at time *k*, the conditional model is

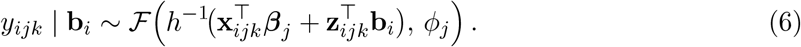

To capture heterogeneity, the random effects are modeled via a finite mixture distribution:

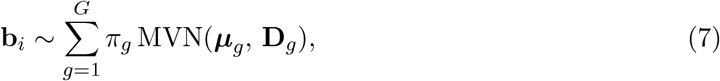

where *π*_*g*_ are the mixing proportions and (***µ***_*g*_, **D**_*g*_) denote the cluster-specific mean vector and covariance matrix.

Posterior inference is performed in the Bayesian framework using MCMC algorithms. Each subject is assigned to the cluster *g* that maximizes the posterior probability Pr(*C*_*i*_ = *g* | **Y**_*i*_), which can again be derived from the mixture distribution of the random effects from equations (6) and (7). BMM can be implemented using the `mixAK` package. In our study, we applied the BMM under Gaussian assumptions with a random slope for time.

### 2.4 Multivariate Longitudinal *k*-Means

The multivariate longitudinal *k*-means method (KML3D) is a nonparametric clustering approach that partitions subjects into *G* non-overlapping clusters by minimizing within-cluster variability [25]. Unlike model-based approaches, KML3D does not assume any specific distributional form and produces hard cluster assignments rather than probabilistic memberships.

Implemented in the `kml3d R` package, the algorithm treats each subject’s multivariate longitudinal profile as a matrix and assigns individuals to the cluster with the closest mean trajectory in terms of Frobenius distance. For subject *i*, the multivariate trajectory is represented as a

*p* × *t* matrix **Y**_*i*_ = (*y*_*ijk*_)_*j*=1,…,*p*; *k*=1,…,*t*_, where *p* is the number of variables and *t* is the number of time points. The clustering objective is to solve

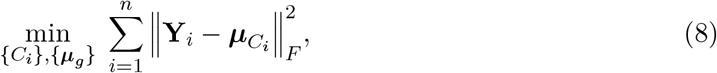

where ∥·∥_*F*_ denotes the Frobenius norm. The cluster centroid ***µ***_*g*_ represents the mean trajectory pattern for cluster *g*:

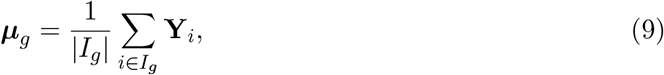

with *I*_*g*_ denoting the set of individuals assigned to cluster *g*.

## 3 Simulation, imputation and evaluation

### 3.1 Data structure

We considered three scenarios to reflect distinct real-world longitudinal data structures.

i. **Scenario 1 (SCE1)** involved metagenomic data with *p* = 80 microbial taxa measured across *t* = 3 and *t* = 5 visits. This dataset was fully observed with no missing values.
ii. **Scenario 2 (SCE2)** consists of data from a repeated-measures multisite study [32] with *p* = 2 clinical measures: composite scores for anxiety and depression, as measured by the Montgomery–Asberg Depression Rating Scale (MADRS) [33] and the Hamilton Anxiety Rating Scale (HARS) [34], respectively, collected over *t* = 14 visits across 24 months.
iii. **Scenario 3 (SCE3)** involved comprehensive behavioral assessment data originating from the same study as SCE2 [32], with more detailed *p* = 24 item-level scores across the same *t* = 14 visits.

### 3.2 Missing data and imputation methods

Both SCE2 and SCE3 exhibited substantial missingness (15–40% across time points). Due to the incompatibility with missing data for most trajectory-clustering methods, we address missingness by evaluating five imputation strategies:

i. **Complete Case (CC):** By retaining a subset of subjects with complete records across all *t* = 14 visits. We recognize that this conservative approach has been the mainstream solution in domain research where incomplete cases are dropped before clustering; however, this approach induces bias if patterns of missingness are not completely at random (or MCAR) [15].
ii. **MICE-Cross-sectional (MICE-CS):** MICE, i.e., multiple imputation by chained equations [35], using demographic covariates (age, sex, race, ethnicity) and other variables measured within the same visit; this imputation strategy retains the between-subject heterogeneity.
iii. **MICE-Longitudinal (MICE-L):** Using predictive mean matching in MICE, we imputed each feature using each person’s own trajectory across time points to exploit within-subject temporal correlation.
iv. **LME-Selective (LME-S):** Model-based imputation using linear mixed-effects models (LME) with subject-specific random intercepts and time-varying fixed effects. In this hybrid strategy, only missing values are replaced by LME predictions, while all observed data are retained.
v. **LME-Full (LME-F):** Model-based replacement of all entries using LME with subject-specific random intercepts and time-varying fixed effects, such that both missing and observed values are replaced by model-based predictions here.

### 3.3 Resampling using eCDF-Copula

To more closely approximate the complexity of real-world longitudinal data, we develop a new simulation strategy based on stratified resampling of observed datasets following imputation. The objective is to construct balanced, homogeneous resampled groups that reliably produce a prespecified number of clusters for downstream evaluation. To achieve this, we extend the nonparametric empirical distribution function (eCDF)-copula simulation framework [36] to longitudinal settings, which preserves both the marginal feature distributions and the empirical dependence structures. This extension flexibly accommodates non-Gaussian and multi-modal distributions, while also maintaining within-visit (cross-feature) correlations and between-visit (within-subject) temporal dependencies. The resampling procedure is detailed in Algorithm 1.

#### Algorithm 1

Longitudinal data resampling framework via eCDF-Copula

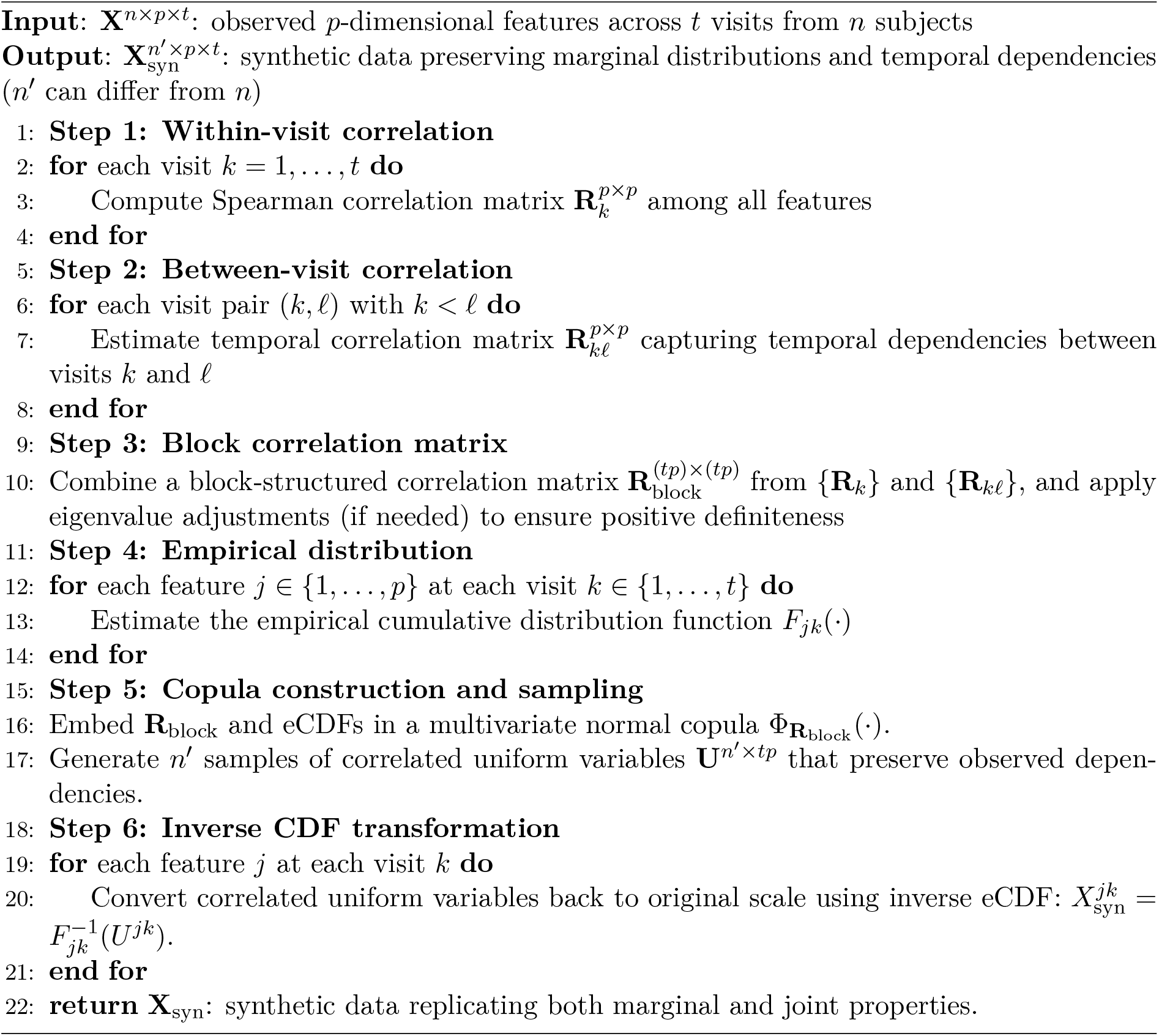

For simulation conditions SCE2 and SCE3, each imputed dataset was stratified and resampled to yield balanced datasets comprising *n* = 200 subjects per group (two groups in total). The resampling preserved key empirical properties of the original data, including marginal distributions, within-subject correlations (intraclass correlation coefficients ranging from 0.4 to 0.6), and cross-feature dependence structures.

For SCE1, which involved fully observed data, resampling was conducted directly to generate three balanced groups without the imputation step.

### 3.4 Clustering approaches

We benchmarked the five longitudinal clustering algorithms described above. Specifically, the group-based trajectory modeling (GBTM) approach, implemented in the `gbmt` and `flexmix` packages, which fits finite mixtures of longitudinal regression models via maximum likelihood. The `lcmm` package extends this framework by incorporating mixed-effects specifications, flexible link functions, and random effects. The `mixAK` package implements the Bayesian mixture model (BMM), estimating multivariate normal mixture components through MCMC sampling. Finally, `kml3d` applies a nonparametric *k*-means algorithm to partition subjects based on multivariate longitudinal trajectories.

Although both `gbmt` and `flexmix` belong to the GBTM family, they differ in flexibility and usability. The `gbmt` package is primarily for continuous time variables and models individual trajectories directly. It is easy to implement, robust to missing values, and performs well with minimal tuning. In contrast, `flexmix` offers greater modeling flexibility, accommodating both continuous and categorical representations of time (e.g., treating time as a factor). However, it does not support missing data and generally requires more careful model specification.

Regarding modeling assumptions, `gbmt`, `flexmix`, and `lcmm` are semi-parametric approaches, combining finite mixture modeling with flexible polynomial or link functions. The BMM implemented in `mixAK` is fully parametric, assuming multivariate normal mixture distributions. In contrast, `kml3d` is fully nonparametric, assigning subjects to hard clusters by minimizing within-cluster distances without any explicit distributional assumptions.

For SCE3 with *p* = 24 item-level scores, we applied principal component analysis (PCA) to reduce dimensionality prior to clustering, which is consistent with common practice in high-dimensional, high-timepoint datasets [37].

### 3.5 Evaluation criteria

#### Adjusted Rand Index (ARI)

We chose the widely used external criterion ARI for cluster validation. It is based on pairwise comparisons of subjects: two subjects are considered concordant if they are assigned to the same cluster in both partitions, or to different clusters in both partitions. The ARI takes values in [™1, 1], with 1 indicating perfect agreement, 0 corresponding to the expected value under random clustering, and negative values indicating worse-than-random agreement [14].

The ARI quantifies the similarity between two partitions by adjusting for chance agreement, which corrects the Rand Index by subtracting the expected similarity between two random clusterings [38, 39]:

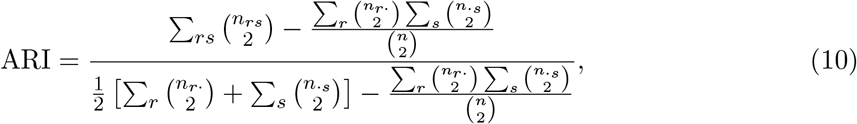

where *n*_*rs*_ is the number of subjects simultaneously assigned to cluster *r* in the true partition and cluster *s* in the estimated partition, *n*_*r*_ and *n* _*s*_ denote row and column sums of the contingency table, and *n* is the total number of subjects.

#### Calinski-Harabasz Index (CHI)

We used CHI as an internal criterion that evaluates the ratio of between-cluster dispersion to within-cluster dispersion [40]. For a partition into *K* clusters, it is defined as

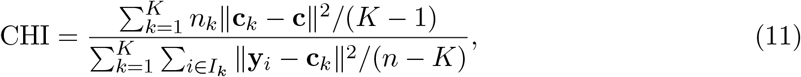

where *n*_*k*_ is the size of cluster *k*, **c**_*k*_ is its centroid, **c** is the overall centroid, **y**_*i*_ is the vectorized trajectory matrix vec(**Y**_*i*_) of subject *i*, and *n* is the total sample size. Higher values indicate better separation between clusters and stronger within-cluster homogeneity [41]. The index is non-negative, with no fixed upper bound.

To evaluate clustering performance, we considered two situations. When the number of clusters was known (i.e., data simulated with a prespecified group structure), we assessed the clustering validity using the Adjusted Rand Index (ARI) with the true labels. When the number of clusters was unknown, we first selected the optimal number of clusters according to the Calinski-Harabasz Index (CHI) using candidate values *K* = 2, …, 5, and then evaluated the selected solution using ARI.

## 4 Results

### 4.1 Performance of clustering methods across scenarios

#### 4.1.1 Clustering Accuracy

In SCE1 (high-dimensional/low-timepoint, see Figure 1), `gbmt`, followed by `kml3d`, are the best clustering methods. As another GBTM-based approach, `flexmix` showed unstable performance. It is competitive under the *t* = 5 setting but less promising when *t* = 3.

**Figure 1.**
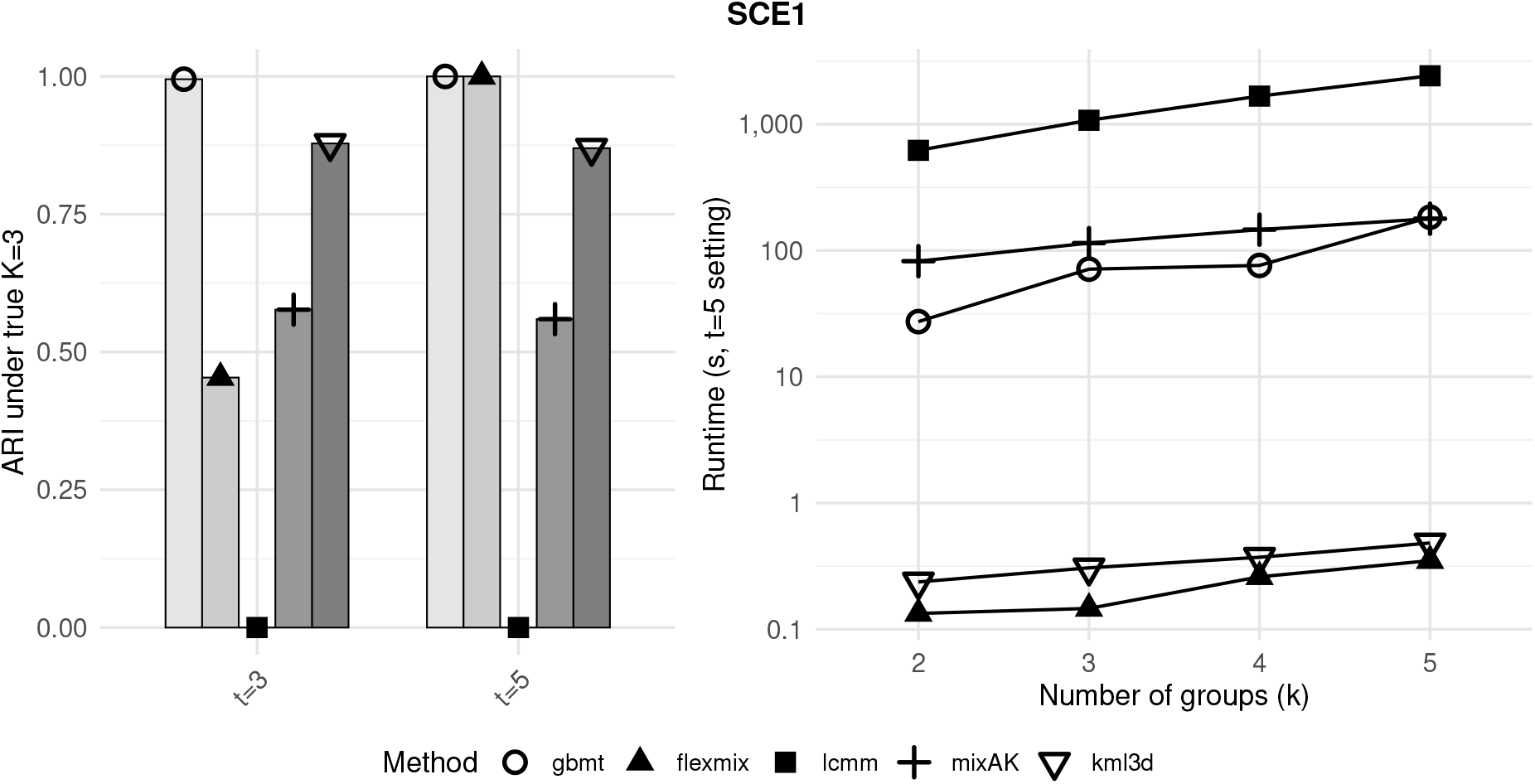
Clustering performance of different methods under Scenario 1 (SCE1). Left: ARI values at the true number of clusters (*K* = 3) under the *t* = 3 and *t* = 5 settings. Right: Runtime (s) across different numbers of clusters (*K* = 2–5), under the *t* = 5 setting.

`gbmt` also outperformed the other methods in SCE2 and SCE3. In SCE2 (low-dimensional/ high-timepoint, see Figure 2), `flexmix` performed relatively well and was consistently ranked second to `gbmt` across different imputation methods (CC, MICE-CS, MICE-L, LME-S, LME-F). However, in SCE3 (high-dimensional/high-timepoint, see Figure 2), `flexmix` performance deteriorated substantially. Interestingly, `lcmm`, which is based on mixed-effects models, often failed to converge due to the increased model complexity, particularly in SCE3. Moreover, when convergence was achieved, nearly all samples were trivially assigned to a single class, regardless of the specified number of clusters (*K* ≠ 1). This likely reflects that the fixed-effect specification already assigns each subject a unique random intercept, leaving insufficient residual structure for meaningful clustering [42].

**Figure 2.**
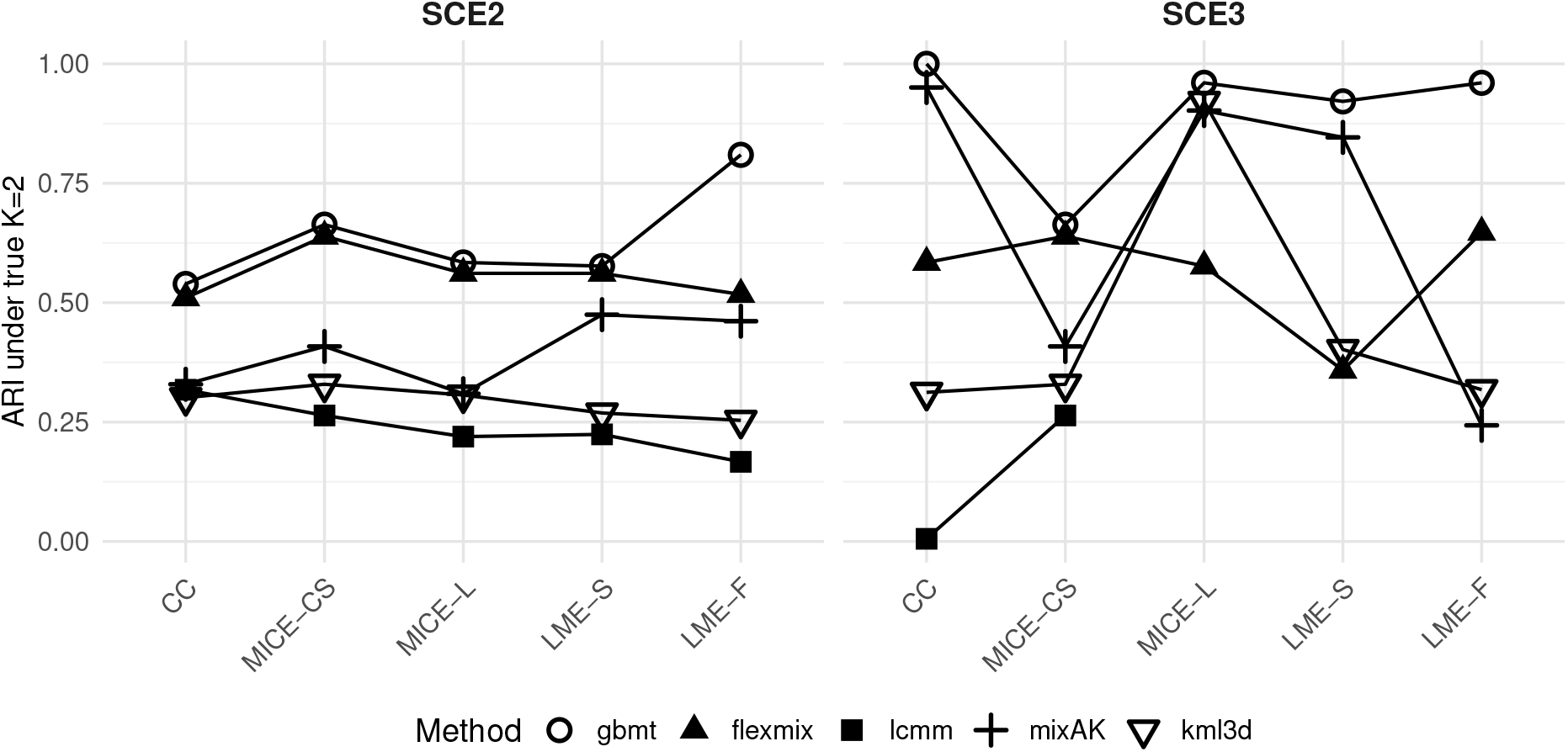
Clustering performance of different methods under Scenario 2 (SCE2) and Scenario 3 (SCE3). Adjusted Rand Index (ARI) values are shown at the true number of clusters (*K* = 2) across different imputation strategies: CC (Complete Case), MICE-CS (MICE–Cross-sectional), MICE-L (MICE–Longitudinal), LME-S (LME–Selective), and LME-F (LME–Full). Missing `lcmm` points indicate settings in which the model failed to converge.

Across all three scenarios and under all settings, `gbmt` consistently achieved the highest accuracy when the true number of clusters was known or specified. In contrast, `lcmm` performed the worst across settings, driven by our observation that it frequently assigns all subjects into a single cluster regardless of *K*. `kml3d`, `flexmix` and `mixAK` only showed competitive performance under certain scenarios and specific imputation strategies. Specifically, `kml3d` performed well in SCE1 and under MICE-L in SCE3, `flexmix` showed strong performance in SCE2, and `mixAK` was competitive under CC, MICE-L, and LME-S in SCE3.

### 4.1.2 Runtime Comparison

As shown in Figure 1, `flexmix` and `kml3d` were the most computationally efficient, whereas `gbmt`, `lcmm`, and `mixAK` required much longer runtimes (e.g., 0.5 vs. 1000 seconds), with `lcmm` and `mixAK` frequently encountering convergence failures in high-dimensional settings. The `mixAK` generally achieved moderate accuracy (ARI: 0.3–0.6 in SCE1 and SCE2) across scenarios but was limited by extremely long runtimes and complex implementation requirements.

In summary, `gbmt` appeared to be the most robust and accurate method across diverse longitudinal settings, while `flexmix` and `kml3d` are fast but their performances are scenario-dependent.

### 4.2 Impact of imputation strategies

The motivation for imputing missing values prior to clustering stems from the realities of longitudinal biomedical datasets, where incomplete repeated measures are inevitable. Evaluating whether clustering performance remains robust across multiple imputation strategies is thus essential. This robustness was particularly evident in SCE2 when feature dimension is low (see Figure 2, left panel), where performance was largely consistent across imputation methods.

It is well recognized that different imputation strategies yield varying levels of performance depending on the dataset characteristics and missing patterns [43]. Our findings show that no single imputation method universally outperforms across all scenarios or data structures.

Nonetheless, several noteworthy patterns emerged from our simulations. In SCE2, `gbmt` combined with LME-F imputation achieved the most distinct subtype separation and the highest ARI. In contrast, in SCE3, the choice of imputation method had a much stronger influence on clustering outcomes (e.g., for `mixAK`: ARI = 0.25 under LME-F vs. 0.80 under MICE-L), emphasizing the importance of selecting an appropriate imputation strategy.

PCA projections (Figure 3) further illustrated how imputation alters the data structure. In SCE2, only LME-F yielded well-separated groups (Control vs. Remitted) in the first two PCs, while other methods yielded overlapping clusters. This improved separation translated directly into superior clustering performance of `gbmt` (ARI = 0.81). By contrast, SCE3 demonstrated clear group separation under all five imputations, and clustering accuracy in SCE3 (high feature dimension) generally exceeded that in SCE2 (low feature dimension) under comparable settings. Hence, an effective practice for applied researchers may be to evaluate multiple imputation strategies, visualize the dimension-reduction plots, and select methods that preserve apparent subtype separation.

**Figure 3.**
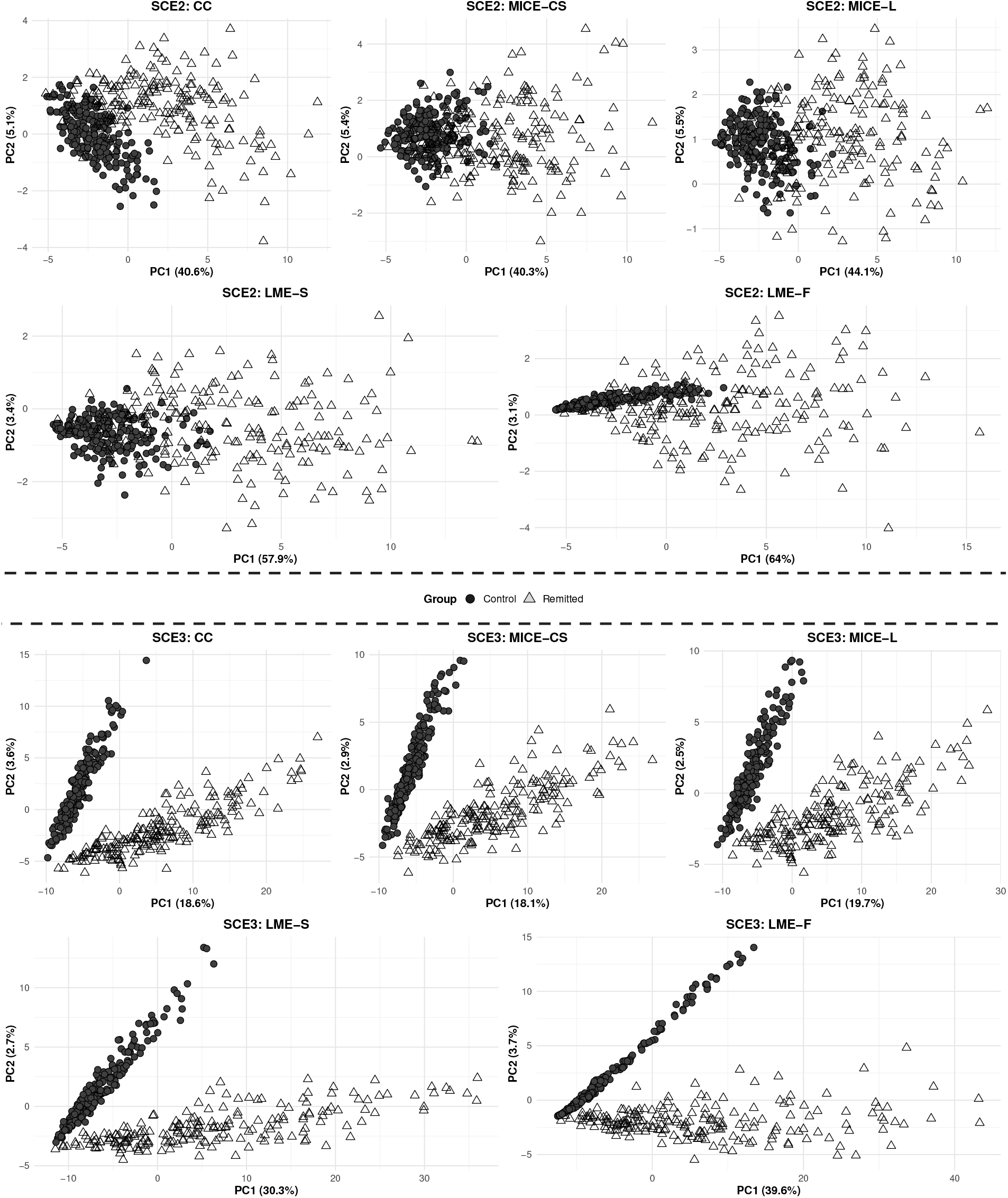
PCA projection scatterplots on the PC1–PC2 plane for Scenario 2 (SCE2, upper panel) and Scenario 3 (SCE3, lower panel) after applying different imputation strategies: CC (Complete Case), MICE-CS (MICE–Cross-sectional), MICE-L (MICE–Longitudinal), LME-S (LME–Selective), and LME-F (LME–Full).

For the overall clustering accuracy, MICE-CS consistently underperformed relative to its longitudinal counterpart, MICE-L, especially when paired with `gbmt`. This discrepancy reflects MICE-CS’s reliance on within-visit correlations, which fail to capture temporal dependencies that MICE-L leverages more effectively. Complete-case (CC) analysis, though appealing for its simplicity, assumes data are missing completely at random (MCAR), an assumption that may not hold in real-world longitudinal studies.

As expected, clustering performance was generally stronger in high-dimension (SCE3) than in low-dimension settings, given that SCE3 included 24 features compared to only 2 in SCE2, thereby offering richer information for subgroup discrimination. The PCA visualizations confirmed this, showing markedly clearer separation in SCE3. Certain algorithms such as `mixAK` also exhibited better performances under selected imputations (CC, MICE-L, and LME-S) in SCE3, though their overall stability remained limited.

In summary, the impact of the imputation strategy on clustering performance depends strongly on the underlying data structure. These results emphasize that imputation is not just a preprocessing step but an integral modeling decision that can substantially alter clustering outcomes. Accordingly, adopting different imputation strategies, followed by diagnostic visualization and comparative evaluation, will be recommended to ensure robust and interpretable results.

### 4.3 When *K* is unknown: how to determine the number of clusters

Determining the optimal number of clusters (*K*) remains a critical yet challenging aspect of longitudinal clustering, especially in high-dimensional settings [15]. Among several candidate criteria, we selected the Calinski–Harabasz Index (CHI) to determine *K* because this distance-based metric is broadly applicable across all five clustering algorithms considered. This choice ensures fairness in benchmarking, as likelihood-based measures (e.g., BIC or AIC) are only defined for model-based approaches. After selecting *K*, clustering performance was evaluated using the Adjusted Rand Index (ARI) to quantify agreement with the true group labels.

When *K* was unknown and had to be estimated, the previous poorly performing algorithms, such as `lcmm`, tended to yield biased estimates of *K*. However, the ability of otherwise strong methods such as `gbmt` to recover the true number of clusters was substantially impaired under LME-based imputations, particularly in SCE3, resulting in performance declines (Figure 4).

**Figure 4.**
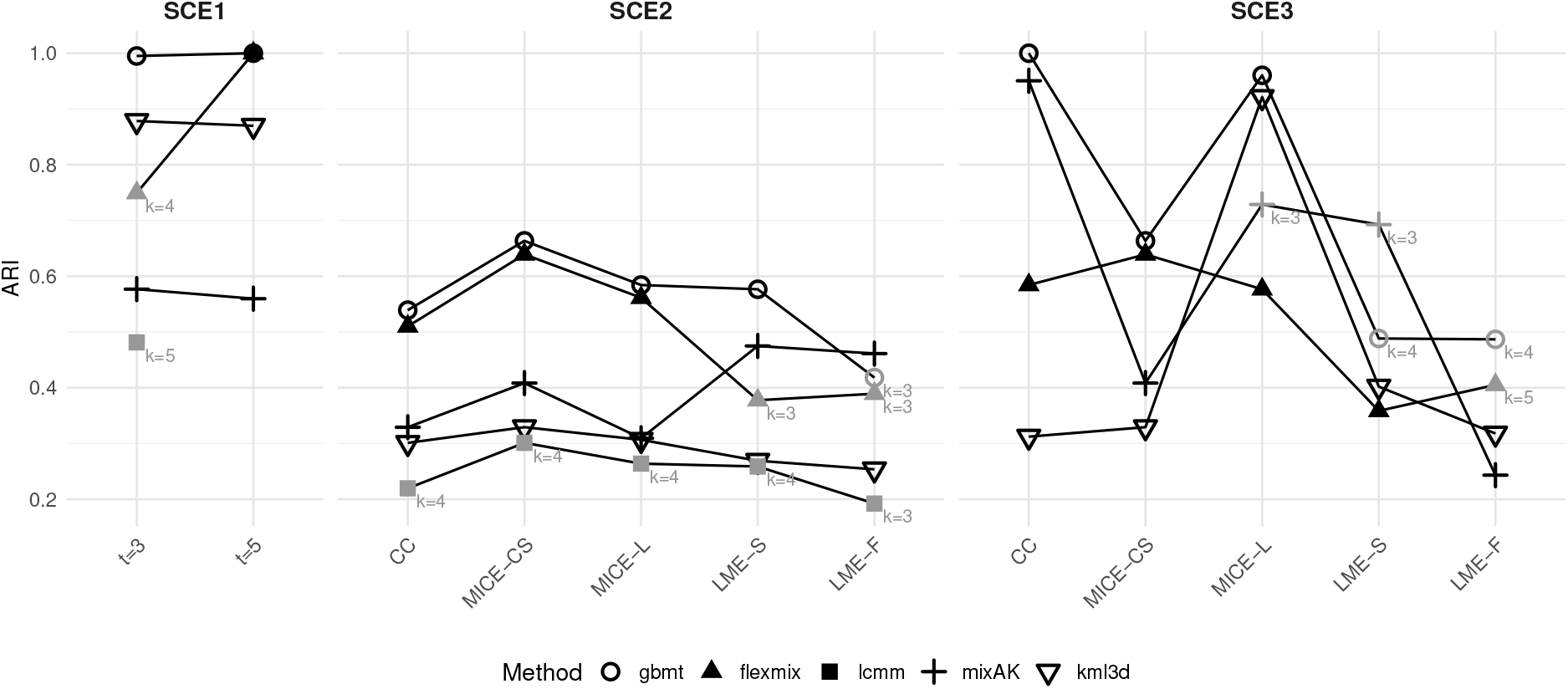
Adjusted Rand Index (ARI) values at the best number of clusters (*K*) determined by the Calinski–Harabasz Index (CHI). Results are shown for Scenario 1 (SCE1) under the *t* = 3 and *t* = 5 settings, and for Scenario 2 (SCE2) and Scenario 3 (SCE3) under different imputation strategies: CC (Complete Case), MICE-CS (MICE–Cross-sectional), MICE-L (MICE–Longitudinal), LME-S (LME–Selective), and LME-F (LME–Full). Black indicates cases where the CHI-selected *K* matched the true number of clusters, while gray indicates incorrect estimates (with the estimated *K* labeled beside the point). Missing `lcmm` points indicate settings in which the model failed to converge.

For imputation methods, LME-S and LME-F in SCE3 resulted in notably poorer performance. As illustrated in Figure 3 (lower panel), these imputations, particularly LME-F, produced strongly distorted distributions in the PCA-reduced space. Consistently, Table 1 shows that CHI values computed at the true labels were substantially lower for these LME-based imputations, indicating that distance-based indices such as CHI may fail to capture the true subgroup once the data structure is distorted by model-based imputations. Furthermore, in SCE3, applying `gbmt` to the LME-F–imputed dataset with true *K* = 2 yielded an almost perfect clustering (ARI ≈ 1.0); however, the CHI for this correct solution (166.90) was lower than that of a more biased clustering solution (CHI = 207.93), showing a clear disconnect between CHI magnitude and true cluster recovery.

**Table 1:** Calinski–Harabasz Index (CHI) evaluated on the ground-truth two-cluster partition after applying five imputation strategies in SCE2 and SCE3: CC (Complete Case), MICE-CS (MICE–Cross-sectional), MICE-L (MICE–Longitudinal), LME-S (LME–Selective), and LME-F (LME–Full). In SCE3, LME-based imputations (LME-F, LME-S) yield markedly lower CHI than non-LME methods, indicating a reduced between-group separation, which can bias CHI-based selection of *K*.

| Scenario | Method | CHI (true labels) |
| --- | --- | --- |
| SCE2 | CC | 181.23 |
| SCE2 | MICE-CS | 192.71 |
| SCE2 | MICE-L | 222.59 |
| SCE2 | LME-S | 260.97 |
| SCE2 | LME-F | 255.11 |
| SCE3 | CC | 265.09 |
| SCE3 | MICE-CS | 272.27 |
| SCE3 | MICE-L | 262.03 |
| SCE3 | LME-S | **199.35** |
| SCE3 | LME-F | **152.80** |

Methods based on group-based trajectory modeling, such as `gbmt` and `flexmix`, appeared particularly sensitive to this distortion caused by LME-based imputations. In SCE3, the most severe drop in ARI was observed for `gbmt`, decreasing from nearly 1 to around 0.53 under LME imputations. A similar but milder decline was also observed for `mixAK`, where ARI decreased from approximately 0.85 to 0.70.

Taken together, these results demonstrate that distance-based criteria such as CHI are vulnerable to bias following LME-based imputations, which can distort the data structure. Caution is therefore warranted when using CHI or similar metrics to estimate *K* in such settings. More robust alternatives, such as ensemble approaches that aggregate multiple indices [44], may provide more reliable guidance for determining the optimal number of clusters in future applications.

### 4.4 Validating fidelity of eCDF-copula resampling

Lastly, we validated whether the adopted eCDF-copula framework faithfully reproduced the global structure, correlation patterns, and marginal distributions of the real-world longitudinal data.

First, dimensionality-reduction analyses showed strong concordance between simulated and original samples (Supplementary Fig. 1). For both the Control and Remitted groups, we combined the original and simulated wide-format datasets and embedded them into lower-dimensional spaces (e.g., via UMAP/t-SNE/PCA), labeling points by data source (original vs. simulated). The resulting embeddings allowed us to assess whether the simulated data preserved the intrinsic linear and nonlinear structures present in the original datasets. UMAP achieved 98.0% (Control) and 96.7% (Remitted) overlap, t-SNE 92.9% and 88.7%, and PCA 73.8% and 78.1%, indicating that both linear and nonlinear structures were all well preserved.

At the correlation level, full (*tp*) × (*tp*) matrices revealed that within-visit and between-visit dependencies were accurately replicated. Simulated heatmaps (Supplementary Fig. 2) mirrored the block patterns of the original data, confirming the well-retained temporal correlations.

Marginal distributions were also faithfully reproduced: overlaid eCDF curves showed near-perfect alignment between simulated and observed values across time points, with preserved skewness, multimodality, and bounded ranges (Supplementary Fig. 3).

Together, these results validate that the eCDF-copula approach generates synthetic longitudinal datasets that retain the complex multivariate structure of real data, thereby supporting its use for methodological evaluation in our analyses.

## 5 Real-World Application

The clustering methods evaluated above demonstrate the advantage of the `gbmt` package across different scenarios and settings. To further exemplify, we applied four clustering methods, `gbmt`, `flexmix`, `lcmm`, and `mixAK`, on a real clinical dataset from the Chronic Hypertension and Pregnancy (CHAP) project [45]. This study followed women longitudinally from randomization until delivery, with repeated measurements of blood pressure throughout pregnancy. Since not all subjects have the same number of longitudinal observations, our illustration did not include the `kml3d` method.

We analyzed the Period 2 of the dataset, which contains longitudinal follow-up data from 3,934 pregnant participants. For each participant, multiple repeated blood pressure measurements were recorded between 3 and 40 weeks of gestation. At each visit, systolic blood pressure (SBP) and diastolic blood pressure (DBP) were collected. The dataset also includes demographic and health-related information such as age, body mass index (BMI), race, diabetes, chronic kidney disease (CKD), smoking status, and parity (e.g., nulliparous status).

To prepare data for clustering, we applied a two-stage dimension reduction approach. First, functional principal component analysis (FPCA) was applied separately to SBP and DBP trajectories using the Fast Covariance Estimation (FACE) algorithm [46, 47], which handles irregular and sparse longitudinal data by reconstructing smooth functional curves. For each blood pressure type, components explaining 99% of variance were retained, yielding 4 fPCs for SBP and 4 fPCs for DBP. These 8 fPC scores were then combined into a unified feature matrix. To address potential correlations between SBP and DBP features, we applied conventional principal component analysis (PCA) to the combined fPC scores. The top 5 principal components were retained, which cumulatively explained 88.5% of the total variance. Since the true *K* is unknown in practice, for each method, we evaluated clustering solutions across *K* = 2, …, 5 using the Calinski–Harabasz Index (CHI), and selected the *K* corresponding to the maximum CHI (Table 2). `flexmix`, `gbmt`, and `lcmm` all estimated *K* = 2, whereas only `mixAK` selected *K* = 3. A comparison of the clustering assignments projected onto the first two principal components is shown in Figure 5.

**Table 2:** Calinski–Harabasz Index (CHI) values for clustering solutions across different methods and numbers of clusters (*K*). The maximum CHI for each method is shown in bold.

| Method | $K$ | CHI |
| --- | --- | --- |
| flexmix | **2** | **571.66** |
| flexmix | 3 | 379.62 |
| flexmix | 4 | 278.37 |
| flexmix | 5 | 216.33 |
| gbmt | **2** | **488.60** |
| gbmt | 3 | 314.56 |
| gbmt | 4 | 216.96 |
| gbmt | 5 | 168.60 |
| lcmm | **2** | **573.87** |
| lcmm | 3 | 372.47 |
| lcmm | 4 | 449.25 |
| lcmm | 5 | 486.12 |
| mixAK | 2 | 133.04 |
| mixAK | **3** | **242.90** |
| mixAK | 4 | 97.42 |
| mixAK | 5 | 91.14 |

**Figure 5.**
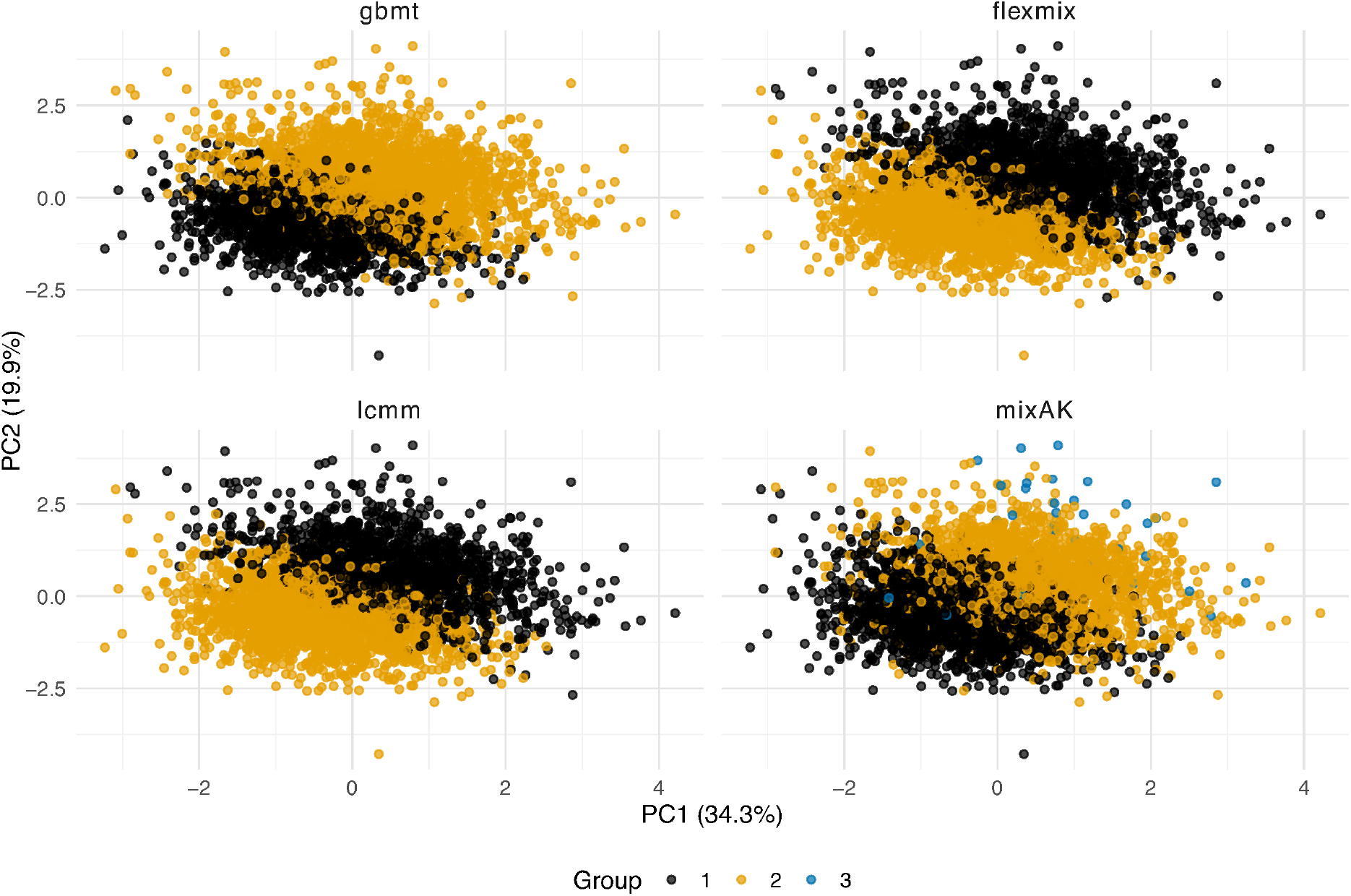
Clustering results of the CHAP Period 2 dataset projected onto the first two principal components (PC1: 34.3%, PC2: 19.9%). `gbmt`, `flexmix`, `lcmm`, and `mixAK` were applied to the longitudinal blood pressure data after FPCA–PCA dimensionality reduction. For `gbmt`, `flexmix`, and `lcmm`, the Calinski–Harabasz Index indicated an optimal solution with *K* = 2 clusters, separating participants into lower-risk (black) and higher-risk (ochre) blood pressure groups. `mixAK` selected *K* = 3, identifying an additional small subgroup (blue, *n* = 40) while maintaining consistency with the two-group structure.

The results indicate that all four methods yielded broadly consistent clustering structures. Given that this dataset reflects longitudinal blood pressure trajectories in a population with potential risk for chronic hypertension, an effective clustering method would be expected to separate participants with persistently elevated blood pressure from those with lower levels. As shown in Figure 5, the two-group solutions from `gbmt`, `flexmix`, and `lcmm` consistently partitioned participants into lower-risk and higher-risk blood pressure groups. Although `mixAK` produced three clusters, the third group contained only 40 participants and was relatively small. Further, the spaghetti plots based on the `gbmt`-derived clustering assignments (Figure 6) clearly distinguished participants exhibiting lower vs. higher average trajectories of both SBP and DBP across pregnancy. These findings suggest that the clustering results align with clinically meaningful heterogeneity in blood pressure patterns.

**Figure 6.**
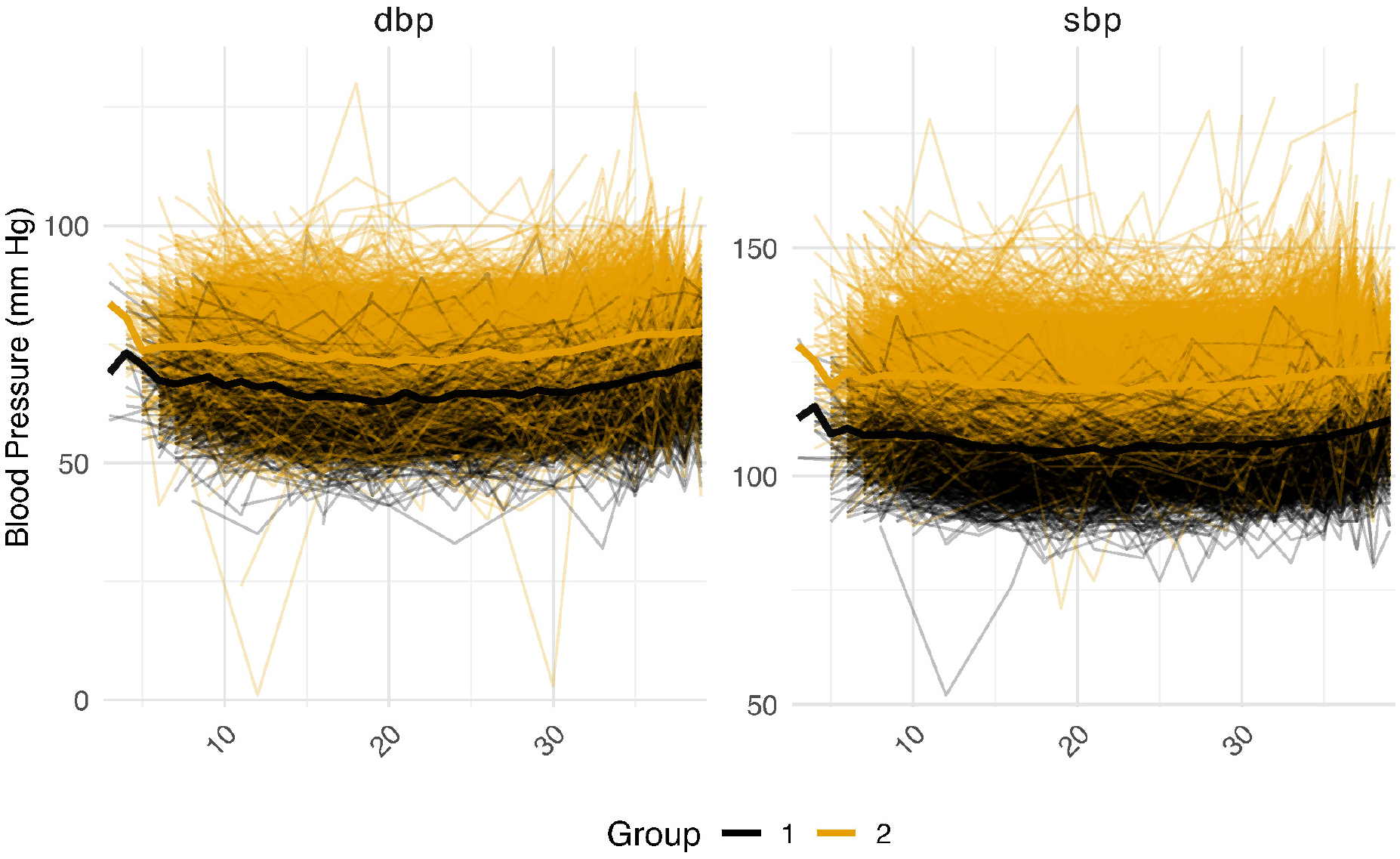
Spaghetti plots of longitudinal blood pressure trajectories after `gbmt` clustering. The left panel shows diastolic blood pressure (DBP) and the right panel shows systolic blood pressure (SBP). Each thin line represents an individual participant’s repeated measurements during pregnancy, color-coded by `gbmt`-assigned cluster (Group 1: black; Group 2: ochre). Thick lines denote the mean trajectory within each cluster.

Although no gold-standard labels are available as in most real-world studies, the consistent findings across clustering methods, combined with the visual coherence of cluster patterns, support their utility in identifying latent subgroups within real-world longitudinal clinical data.

## 6 Discussion

This study systematically evaluated joint clustering methods for multivariate longitudinal trajectories, addressing the challenges of identifying latent subgroups in complex datasets characterized by high dimensionality, temporal correlation, and missing data. While previous studies have primarily focused on single longitudinal features or cross-sectional data, comprehensive evaluations in realistic multivariate longitudinal studies remain scarce. We addressed this gap by benchmarking five representative clustering algorithms across three real-world scenarios and developing a novel eCDF-copula-based resampling framework that accurately reproduces empirical data distributions for robust evaluation of real-world clustering performance. Collectively, these efforts provide both methodological insights and practical guidance for selecting clustering strategies in applied research.

Our first finding was that `gbmt` consistently achieved the highest clustering accuracy across scenarios and imputation settings, confirming its robustness for capturing heterogeneous temporal trajectories. In contrast, `lcmm` frequently returned trivial or unstable solutions in high-dimensional settings, and was less robust overall. `flexmix` and `kml3d` offered computational efficiency but demonstrated scenario-specific sensitivity. These results underscore the trade-off between flexibility and computational scalability among existing approaches.

Our second finding highlights the decisive role of the imputation strategy. In low-dimensional trajectories, clustering performance was relatively stable across methods; however, in high-dimensional settings, performance varied markedly depending on the imputation approach. Temporal methods such as MICE-Longitudinal (MICE-L) outperformed cross-sectional or other strategies, which reflects the importance of preserving within-subject correlations. When missingness is approximately MCAR, a simple complete-case analysis remains acceptable; however, when this assumption is violated, careful comparison across imputation methods becomes essential to ensure robust subgroup recovery.

A third key observation concerns the determination of the number of clusters. Distance-based indices such as the Calinski-Harabasz Index (CHI) tended to misestimate cluster numbers following LME imputation, likely due to the distorted structure introduced by model-based smoothing. For example, in SCE3, `gbmt` under LME-Full imputation achieved an ARI close to 1 with *K* = 2, yet the CHI favored biased alternatives. These findings caution against sole reliance on distance-based criteria and suggest that likelihood-based or ensemble approaches may yield more reliable solutions. The strengths of this study lie in its integration of realistic data generation and comprehensive benchmarking. Our eCDF–copula framework offers a robust, nonparametric mechanism for generating realistic longitudinal datasets that retain observed distributions and dependence structures. By enabling reproducible and flexible benchmarking, it bridges the persistent gap between theoretical method development and the heterogeneity of real-world data. Unlike prior reviews [15] based on simplified simulation schemes, our results show that methodological insights can shift considerably when evaluated under empirically grounded conditions.

Several limitations warrant consideration. First, although our nonparametric resampling enhances realism, its generalizability to datasets with alternative missingness mechanisms (e.g., MAR or MNAR) remains to be examined. Second, further work is needed to refine cluster-number selection, as traditional metrics such as CHI systematically underestimate *K* in complex settings. Addressing these limitations will further strengthen the reliability and applicability of the proposed framework.

In conclusion, our findings establish `gbmt` as the most reliable method for identifying latent heterogeneity in multivariate longitudinal data, provided that imputation strategies are carefully matched to data characteristics. The combination of real-world benchmarking and the new eCDF-copula resampling framework delivers both practical guidance for applied researchers and methodological advances for longitudinal data science. Together, these contributions lay the foundation for more rigorous, data-driven approaches to clustering in complex biomedical and epidemiological studies.

## References

[1] Qianheng Ma. Extensions to Statistical Methods for Analyzing Intensive Multilevel Longitudinal Data Collected by Mobile and Wearable Devices. PhD thesis, University of Chicago, 2021.

[2] James Yang, Jacky Kuo, Wei-Chung Su, Douglas E. Jorenby, Megan E. Piper, and Anne Buu. A new statistical model for longitudinal ecological momentary assessment data on dual use of electronic and combustible cigarettes. The American Journal of Drug and Alcohol Abuse, 48(5):529–537, 2022. doi: 10.1080/00952990.2022.2027955.

[3] Ali R. Taheriyoun, Allen Ross, Abolfazl Safikhani, Damoon Soudbakhsh, and Ali Rahnavard. Longitudinal Omics Data Analysis: A Review on Models, Algorithms, and Tools. arXiv preprint arXiv:2506.11161, 2025.

[4] Rebecca A Deek, Siyuan Ma, James Lewis, and Hongzhe Li. Statistical and computational methods for integrating microbiome, host genomics, and metabolomics data. eLife, 13: e88956, 2024. doi: 10.7554/eLife.88956.

[5] Saritha Kodikara, Susan Ellul, and Kim-Anh Lê Cao. Statistical challenges in longitudinal microbiome data analysis. Briefings in Bioinformatics, 23(4):bbac273, 2022. doi: 10.1093/bib/bbac273.

[6] Cécile Proust-Lima, Mbéry Séne, Jeremy MG Taylor, and Hélène Jacqmin-Gadda. Joint latent class models for longitudinal and time-to-event data: A review. Statistical Methods in Medical Research, 23(1):74–90, 2014. doi: 10.1177/0962280212445839.

[7] Bret Zeldow, James Flory, Alisa Stephens-Shields, Marsha Raebel, and Jason A Roy. Functional clustering methods for longitudinal data with application to electronic health records. Statistical Methods in Medical Research, 30(3):655–670, 2021. doi: 10.1177/0962280220965630.

[8] Maéva Kyheng, Génia Babykina, and Alain Duhamel. Joint Latent Class Models: A Tutorial on Practical Applications in Clinical Research. Statistics in Medicine, 44(8–9): e70047, 2025. doi: 10.1002/sim.70047.

[9] Cécile Proust-Lima, Viviane Philipps, and Benoit Liquet. Estimation of Extended Mixed Models Using Latent Classes and Latent Processes: The R Package lcmm. Journal of Statistical Software, 78:1–56, 2017. doi: 10.18637/jss.v078.i02.

[10] Cecile Proust-Lima, Viviane Philipps, Amadou Diakite, and Benoit Liquet. lcmm: Extended Mixed Models Using Latent Classes and Latent Processes, 2025. R package version 2.2.1.

[11] J. MacQueen. Some methods for classification and analysis of multivariate observations. In Proceedings of the Fifth Berkeley Symposium on Mathematical Statistics and Probability, Volume 1: Statistics, volume 5.1, pages 281–298. University of California Press, 1967.

[12] Karen Leffondré, Michal Abrahamowicz, Armelle Regeasse, Gillian A. Hawker, Elizabeth M Badley, Jane McCusker, and Eric Belzile. Statistical measures were proposed for identifying longitudinal patterns of change in quantitative health indicators. Journal of Clinical Epidemiology, 57(10):1049–1062, 2004. doi: 10.1016/j.jclinepi.2004.02.012.

[13] Christophe Genolini and Bruno Falissard. KmL: K-means for longitudinal data. Computational Statistics, 25(2):317–328, 2010. doi: 10.1007/s00180-009-0178-4.

[14] Peter Verboon and Ron Pat-El. Clustering Longitudinal Data Using R: A Monte Carlo Study. Methodology, 18(2):144–163, 2022. doi: 10.5964/meth.7143.

[15] Zihang Lu, Mojtaba Ahmadiankalati, and Zhiwen Tan. Joint clustering multiple longitudinal features: A comparison of methods and software packages with practical guidance. Statistics in Medicine, 42(29):5513–5540, 2023. doi: 10.1002/sim.9917.

[16] Alessandro Magrini. Assessment of agricultural sustainability in European Union countries: A group-based multivariate trajectory approach. AStA Advances in Statistical Analysis, 106(4):673–703, 2022. doi: 10.1007/s10182-022-00437-9.

[17] Bettina Grün and Friedrich Leisch. Flexmix: Flexible Mixture Modeling, 2025. R package version 2. 3-20.

[18] Bettina Grün and Friedrich Leisch. Fitting finite mixtures of generalized linear regressions in R. Computational Statistics & Data Analysis, 51(11):5247–5252, 2007. doi: 10.1016/j.csda.2006.08.014.

[19] Bettina Grün and Friedrich Leisch. FlexMix Version 2: Finite Mixtures with Concomitant Variables and Varying and Constant Parameters. Journal of Statistical Software, 28:1–35, 2008. doi: 10.18637/jss.v028.i04.

[20] Friedrich Leisch. FlexMix: A General Framework for Finite Mixture Models and Latent Class Regression in R. Journal of Statistical Software, 11:1–18, 2004. doi: 10.18637/jss.v011.i08.

[21] Arnošt Komárek. A new R package for Bayesian estimation of multivariate normal mixtures allowing for selection of the number of components and interval-censored data. Computational Statistics & Data Analysis, 53(12):3932–3947, 2009. doi: 10.1016/j.csda.2009.05.006.

[22] Arnošt Komárek and Lenka Komárková. Capabilities of R Package mixAK for Clustering Based on Multivariate Continuous and Discrete Longitudinal Data. Journal of Statistical Software, 59:1–38, 2014. doi: 10.18637/jss.v059.i12.

[23] Zhiwen Tan, Chang Shen, Padmaja Subbarao, Wendy Lou, and Zihang Lu. A Joint Modeling Approach for Clustering Mixed-Type Multivariate Longitudinal Data: Application to the CHILD Cohort Study. arXiv preprint arXiv:2210.08385, 2023.

[24] Christophe Genolini, Xavier Alacoque, Mariane Sentenac, and Catherine Arnaud. Kml and kml3d: R Packages to Cluster Longitudinal Data. Journal of Statistical Software, 65: 1–34, 2015. doi: 10.18637/jss.v065.i04.

[25] C. Genolini, J. B. Pingault, T. Driss, S. Côté, R. E. Tremblay, F. Vitaro, C. Arnaud, and B. Falissard. KmL3D: A non-parametric algorithm for clustering joint trajectories. Computer Methods and Programs in Biomedicine, 109(1):104–111, 2013. doi: 10.1016/j.cmpb.2012.08.016.

[26] Kostas Stoitsas, Saurabh Bahulikar, Leonie de Munter, Mariska A. C. de Jongh, Maria A. C. Jansen, Merel M. Jung, Marijn van Wingerden, and Katrijn Van Deun. Clustering of trauma patients based on longitudinal data and the application of machine learning to predict recovery. Scientific Reports, 12(1):16990, 2022. doi: 10.1038/s41598-022-21390-2.

[27] Julian Wenzel, Nils Dreschke, Esther Hanssen, Marlene Rosen, Andrej Ilankovic, Joseph Kambeitz, Anne-Kathrin Fett, and Lana Kambeitz-Ilankovic. Ecological momentary assessment (EMA) combined with unsupervised machine learning shows sensitivity to identify individuals in potential need for psychiatric assessment. European Archives of Psychiatry and Clinical Neuroscience, 274(7):1639–1649, 2024. doi: 10.1007/s00406-023-01668-w.

[28] Daniel S Nagin, Bobby L Jones, Valéria Lima Passos, and Richard E Tremblay. Group-based multi-trajectory modeling. Statistical Methods in Medical Research, 27(7):2015–2023, 2018. doi: 10.1177/0962280216673085.

[29] Daniel S. Nagin. Analyzing developmental trajectories: A semiparametric, group-based approach. Psychological Methods, 4(2):139–157, 1999. doi: 10.1037/1082-989X.4.2.139.

[30] Garrett M. Fitzmaurice, Nan M. Laird, and James H. Ware. Applied Longitudinal Analysis. Wiley, second edition, 2011.

[31] Arnošt Komárek and Lenka Komárková. Clustering for multivariate continuous and discrete longitudinal data. The Annals of Applied Statistics, 7(1), 2013. doi: 10.1214/12-AOAS580.

[32] Warren D. Taylor, Olusola Ajilore, Helmet T. Karim, Meryl A. Butters, Robert Krafty, Brian D. Boyd, Layla Banihashemi, Sarah M. Szymkowicz, Claire Ryan, Jason Hassenstab, Bennett A. Landman, and Carmen Andreescu. Assessing depression recurrence, cognitive burden, and neurobiological homeostasis in late life: Design and rationale of the REMBRANDT Study. Journal of Mood and Anxiety Disorders, 5:100038, 2024. doi: 10.1016/j.xjmad.2023.100038.

[33] Stuart A. Montgomery and Marie Åsberg. A New Depression Scale Designed to be Sensitive to Change. The British Journal of Psychiatry, 134(4):382–389, 1979. doi: 10.1192/bjp.134.4.382.

[34] Max Hamilton. The Assessment of Anxiety States by Rating. British Journal of Medical Psychology, 32(1):50–55, 1959. doi: 10.1111/j.2044-8341.1959.tb00467.x.

[35] Melissa J. Azur, Elizabeth A. Stuart, Constantine Frangakis, and Philip J. Leaf. Multiple imputation by chained equations: What is it and how does it work? International Journal of Methods in Psychiatric Research, 20(1):40–49, 2011. doi: 10.1002/mpr.329.

[36] J. Liu, Xinlian Zhang, T. Chen, T. Wu, T. Lin, L. Jiang, S. Lang, L. Liu, L. Natarajan, J. X. Tu, T. Kosciolek, J. Morton, T. T. Nguyen, B. Schnabl, R. Knight, C. Feng, Y. Zhong, and X. M. Tu. A semiparametric model for between-subject attributes: Applications to betadiversity of microbiome data. Biometrics, 78(3):950–962, 2022. doi: 10.1111/biom.13487.

[37] Ian T. Jolliffe and Jorge Cadima. Principal component analysis: A review and recent developments. Philosophical Transactions of the Royal Society A: Mathematical, Physical and Engineering Sciences, 374(2065):20150202, 2016. doi: 10.1098/rsta.2015.0202.

[38] William M. Rand. Objective Criteria for the Evaluation of Clustering Methods. Journal of the American Statistical Association, 66(336):846–850, 1971. doi: 10.1080/01621459.1971.10482356.

[39] Lawrence Hubert and Phipps Arabie. Comparing partitions. Journal of Classification, 2 (1):193–218, 1985. doi: 10.1007/BF01908075.

[40] T. Caliński and J. Harabasz. A dendrite method for cluster analysis. Communications in Statistics, 3(1):1–27, 1974. doi: 10.1080/03610927408827101.

[41] Yanchi Liu, Zhongmou Li, Hui Xiong, Xuedong Gao, and Junjie Wu. Understanding of Internal Clustering Validation Measures. In 2010 IEEE International Conference on Data Mining, pages 911–916, 2010. doi: 10.1109/ICDM.2010.35.

[42] Daniel McNeish and Jeffrey R. Harring. Improving convergence in growth mixture models without covariance structure constraints. Statistical Methods in Medical Research, 30(4): 994–1012, 2021. doi: 10.1177/0962280220981747.

[43] Mina Jahangiri, Anoshirvan Kazemnejad, Keith S. Goldfeld, Maryam S. Daneshpour, Shayan Mostafaei, Davood Khalili, Mohammad Reza Moghadas, and Mahdi Akbarzadeh. A wide range of missing imputation approaches in longitudinal data: A simulation study and real data analysis. BMC Medical Research Methodology, 23:161, 2023. doi: 10.1186/s12874-023-01968-8.

[44] Sunghae Jun. An ensemble method for validation of cluster analysis. ResearchGate preprint, 2011.

[45] Alan T. Tita, Jeff M. Szychowski, Kim Boggess, Lorraine Dugoff, Baha Sibai, Kirsten Lawrence, Brenna L. Hughes, Joseph Bell, Kjersti Aagaard, Rodney K. Edwards, Kelly Gibson, David M. Haas, Lauren Plante, Torri Metz, Brian Casey, Sean Esplin, Sherri Longo, Matthew Hoffman, George R. Saade, Kara K. Hoppe, Janelle Foroutan, Methodius Tuuli, Michelle Y. Owens, Hyagriv N. Simhan, Heather Frey, Todd Rosen, Anna Palatnik, Susan Baker, Phyllis August, Uma M. Reddy, Wendy Kinzler, Emily Su, Iris Krishna, Nicki Nguyen, Mary E. Norton, Daniel Skupski, Yasser Y. El-Sayed, Dotum Ogunyemi, Zorina S. Galis, Lorie Harper, Namasivayam Ambalavanan, Nancy L. Geller, Suzanne Oparil, Gary R. Cutter, and William W. Andrews. Treatment for Mild Chronic Hypertension during Pregnancy. New England Journal of Medicine, 386(19):1781–1792, 2022. doi: 10.1056/NEJMoa2201295.

[46] Luo Xiao, Cai Li, William Checkley, and Ciprian Crainiceanu. Fast covariance estimation for sparse functional data. Statistics and Computing, 28(3):511–522, 2018. doi: 10.1007/s11222-017-9744-8.

[47] Yidong Zhou, Han Chen, Su I Iao, Poorbita Kundu, Hang Zhou, Satarupa Bhattacharjee, Cody Carroll, Yaqing Chen, Xiongtao Dai, Jianing Fan, Alvaro Gajardo, Pantelis Z. Hadjipantelis, Kyunghee Han, Hao Ji, Changbo Zhu, Hans-Georg Müller, and Jane-Ling Wang. fdapace: Functional Data Analysis and Empirical Dynamics, 2024. R package version 0.6.0.

